# QuASAR: Quantitative Allele Specific Analysis of Reads

**DOI:** 10.1101/007492

**Authors:** Chris T. Harvey, Gregory A. Moyerbrailean, Gordon O. Davis, Xiaoquan Wen, Francesca Luca, Roger Pique-Regi

**Affiliations:** Center for Molecular Medicine and Genetics, Wayne State University, 540 E Canfield, Scott Hall, Detroit, MI 48201, USA; Department of Biostatistics, University of Michigan, Ann Arbor, MI

## Abstract

**Motivation:** Expression quantitative trait loci (eQTL) studies have discovered thousands of genetic variants that regulate gene expression, enabling a better understanding of the functional role of non-coding sequences. However, eQTL studies are costly, requiring large sample sizes and genome-wide genotyping of each sample. In contrast, analysis of allele specific expression (ASE) is becoming a popular approach to detect the effect of genetic variation on gene expression, even within a single individual. This is typically achieved by counting the number of RNA-seq reads matching each allele at heterozygous sites and testing the null hypothesis of a 1:1 allelic ratio. In principle, when genotype information is not readily available it could be inferred from the RNA-seq reads directly. However, there are currently no existing methods that jointly infer genotypes and conduct ASE inference, while considering uncertainty in the genotype calls.

**Results:** We present QuASAR, Quantitative Allele Specific Analysis of Reads, a novel statistical learning method for jointly detecting heterozygous genotypes and inferring ASE. The proposed ASE inference step takes into consideration the uncertainty in the genotype calls while including parameters that model base-call errors in sequencing and allelic over-dispersion. We validated our method with experimental data for which high quality genotypes are available. Results for an additional dataset with multiple replicates at different sequencing depths demonstrate that QuASAR is a powerful tool for ASE analysis when genotypes are not available.

**Availability:** http://github.com/piquelab/QuASAR

## 1 Introduction

Quantitative trait loci (QTLs) for molecular cellular phenotypes (as defined by Dermitzakis, 2012), such as gene expression (eQTL) (e.g. Stranger *et al.,* 2007), transcription factor (TF) binding (Kasowski et *al.,* 2010), and DNase I sensitivity (Degner et *al.,* 2012) have begun to provide a better understanding of how genetic variants in regulatory sequences can affect gene expression levels (see also Stranger et *al.,* 2007; Gibbs et *al.,* 2010; Melzer et *al.,* 2008; Gieger et *al.,* 2008). eQTL studies in particular have been successful at identifying genomic regions associated with gene expression in various tissues and conditions (e.g., Maranville et *al.,* 2011; Barreiro et *al.,* 2012; Nica et *al.,* 2011; Smirnov et *al.,* 2009; Dimas et *al.,* 2009; Ding et *al.,* 2010; Grundberg et *al.,* 2011; Lee et *al.,* 2014; Fairfax et *al.,* 2014). While previous studies have shown an enrichment for GWAS hits among regulatory variants in lymphoblastoid cell lines (LCLs) (Nica et *al.,* 2010; Nicolae et *al.,* 2010), a full understanding of the molecular mechanisms underlying GWAS hits requires functional characterization of each variant in the tissue and environmental conditions relevant for the trait under study (e.g. estrogen level for genetic risk to breast cancer Cowper-Sal-lari *et al.,* 2012).

The ongoing GTEx project will significantly increase the number of surveyed tissues for which eQTL data are available and will represent a useful resource to functionally annotate genetic variants. However, the number of cell-types and environments explored are a small subset of the presumably larger number of regulatory variants that mediate specific GxE interactions. eQTL studies are expensive, requiring large sample sizes (*n* > 70) which may be difficult to achieve for tissues that are obtained by surgical procedures or are difficult to culture *in vitro.* Even if biospecimens are readily available at no cost, eQTL studies require large amounts of experimental work to measure genotypes and gene expression levels. As the measurement of gene expression using high-throughput sequencing (RNA-seq) is becoming more popular than microarrays, RNA-seq library preparation is also becoming less expensive ($46 per sample) while costs of sequencing are also very rapidly decreasing (for example, 16M reads per sample would cost $49 using a multiplexing strategy). Additionally, the sequence information provided by RNA-seq can be used to call genotypes (Shah *et al.,* 2009; Duitama *et al.,* 2012; Piskol *et al.,* 2013), detect and quantify isoforms (Trapnell *et al.,* 2010; Katz *et al.,* 2010) and to measure allele specific expression (ASE), if enough sequencing depth is available (Degner *et al.,* 2009; Pastinen, 2010).

ASE approaches currently represent the most effective way to assay the effect of a cis-regulatory variant within a defined cellular environment, while controlling for any trans-acting modifiers of gene expression, such as the genotype at other loci (Pastinen, 2010; Kasowski *et al.,* 2010; McDaniell *et al.,* 2010; Skelly *et al.,* 2011; Cowper-Sal-lari *et al.,* 2012; Reddy *et al.,* 2012; Hasin-Brumshtein *et al.,* 2014; McVicker *et al.,* 2013; Kukurba *et al.,* 2014). As such, ASE studies have greater statistical power to detect genetic effects in cis than a traditional eQTL mapping approach when using a small sample size. Additionally, ASE may also be useful to detect epigenetic imprinting of gene expression if ASE is present but no eQTL is detected (Seoighe *et al.,* 2006; Degner *et al.,* 2009).

In the absence of ASE, the two alleles for a heterozygous genotype at a single nucleotide polymorphism (SNP) in a gene transcript are represented in a 1:1 ratio of RNA-seq reads. To
reject the null hypothesis and infer ASE. it is necessary to first identify heterozygous SNPs with high confidence, and then conduct inference to detect a departure from a 50% allelic ratio. While genotyping and ASE are usually considered two separate problems, miscalling a homozygous SNP as heterozygous is likely to induce an error in rejecting the ASE null hypothesis; thus, we argue that the two problems should addressed together.

**Fig. 1.**
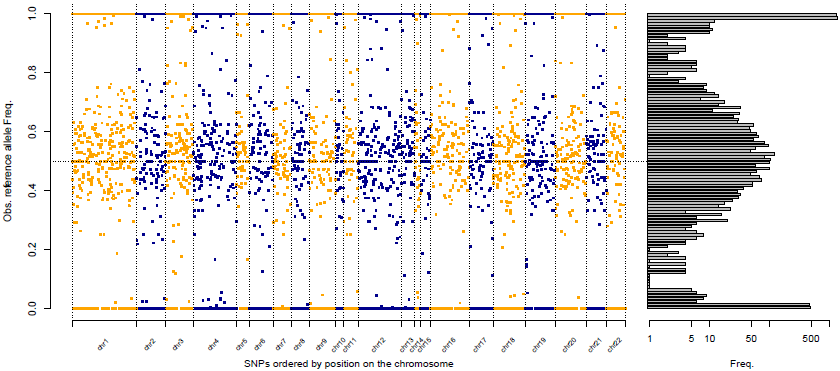
Reference allele frequency from reads overlapping SNPs. (Left) Each dot represents a SNP covered by at least **15** RNA-seq reads. The *y*-axis represents the fraction of RNA-seq reads that match the reference allele (observed 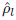). The *x*-axis represents the order of the SNP position in a chromosome. (Right) Histogram showing the distribution of 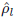 values across the genome. The three modes (*ρ* ∊ {1, 0.5, 0}) correspond respectively to the three possible genotypes: homozygous reference (RR), heterozygous under no ASE (RA), and homozygous alternate (AA).

While it is possible to obtain genotype information from RNA-seq (Shah *et al,* 2009: Duitama *et al*., 2012; Piskol *et al.,* 2013). to the best of our knowledge, all existing methods for detecting ASE consider the genotypes known and error probabilities associated with genotyping are not taken into account for the ASE step. While overall genotyping quality can also be modeled within the ASE model (McVicker *et al.,* 2013). there is currently no method that for each SNP can jointly genotype and detect allelic imbalances in high throughput sequencing data. An approach that takes into account base-calling errors was previously proposed for detecting ASE in ESTs data (Seoighe *et al.,* 2006). but for RNA-seq data it is essential to also include overdispersion. Here, we propose a novel framework for quantitative allele specific analysis of reads (QuASAR) that starts from a single or multiple RNA-seq experiments from the same individual and can directly identify heterozygous SNPs and assess ASE accurately by taking into account base-calling errors and overdispersion in the ASE ratio. QuASAR is evaluated with two different datasets that demonstrate genotyping accuracy and the importance of incorporating the genotype uncertainty when performing ASE inference.

## 2 Methods

### 2.1 QuASAR Approach

QuASAR starts with experimental high-throughput sequencing data. Here we focus on RNA-seq, but the same or similar pipeline can be applied to DNase-seq, ChIP-seq, ATAC-seq or other types of functional genomics library preparations. Figure 1 illustrates the underlying problem: detecting SNPs covered by a number of transcripts with high allelic imbalance, and for which homozygosity (in the presence of base-calling errors) can also be rejected.

We focus our attention on sites that are known to be variable in human populations, specifically we consider all SNPs from the 1000 Genomes project (1KG) with a minor allele frequency MAF > 0.02. We index each SNP with *l* ∊ {1,..., *L*}, and each sample by *s* ∊ {1,..., *S*}. All samples are from the same individual and may represent different experimental conditions or replicates. We only consider SNPs represented in at least 15 reads across all the samples. At each site *l*, three genotypes are possible *gl* ∊ {0,1, 2} being homozygous reference (RR), heterozygous (RA), or homozygous alternate (AA) respectively. For each sample *s* and site *l*, *N_s,l_* represents the total number of reads and 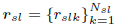 take the value 1 if read *k* matches the reference allele, and 0 if it matches the alternate allele. We can then model the data 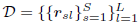 as a mixture model

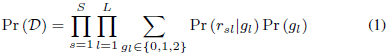

where *Pr(gl)* represents the prior probability associated with each genotype. The probability of the observed reads, 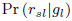, depends on the genotype. For *G_l_* = 0:

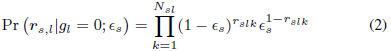

where we will only observe reads matching the alternate allele if those are base-calling errors, here modeled by the parameter *∊_s_*. Conversely, for *G_l_* = 2 we have the following:

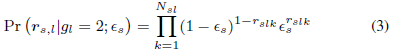

If the genotype is heterozygous *G_l_* = 1, we observe reads from the reference allele with probability *ρl* (or the alternate allele with probability (1 — *ρl*), resulting in the following model:

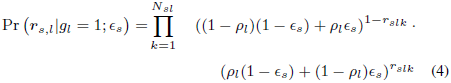

Considering that 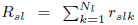 and *A_sl_* = *N_sl_* — *R_sl_* are respectively the number of reads from sample s observed at site *l* matching the reference allele and the alternate allele, the previous equations can be simplified as follows:

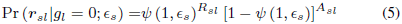

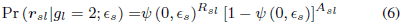

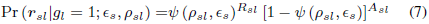

where 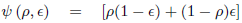 and makes explicit that homozygotes, *gl* = 2 (or *gl* = 0), are indistinguishable from *ρ_ls_* = 0 (or *ρ_ls_* = 1) when *gl* = 1. In QuASAR we resolve this identifiably problem by assuming that those cases with extreme ASE across all replicates are more likely to be homozygous genotypes.

To fit the mixture model we use an EM algorithm (see Methods for more details) in which we estimate sample specific base-calling error rates 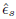 (*ρ* is fixed to 0.5) and posterior probabilities for the genotypes. For the ASE inference step, we wish to test the null hypothesis *ρ_s_l* = 0.5. We additionally consider that Ψ in (5-7) is a random variable *Ψ_sl_* sampled from a ~Beta(*α_sl_*, *β_sl_*) distribution with:

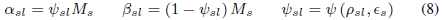

where the hyper parameter *M_s_* controls for over-dispersion and results in a better calibrated test as shown in the Results section. Combining (7) and (8) results in a Beta-binomial distribution (13), which we use to model the number of reads coming from the reference allele. Formalized as a likelihood ratio test (LRT), the inference step takes into account over-dispersion and genotype uncertainty:

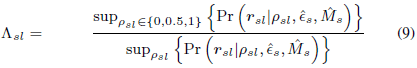

where the set of parameters 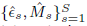 are maximum likelihood estimates (MLE) under the null hypothesis *ρ_ls_* = 0.5 (see Methods section). To calculate a *p*-value we use the property that —2log(*Λ_sl_*) is asymptotically distributed as 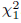.

### 2.2 Model Fitting and Parameter Estimation

To use the expectation maximization (EM) procedure (McLachlan and Krishnan, 2007), we first convert (1) to a “complete” likelihood, as if we knew the underlying genotypes 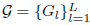:

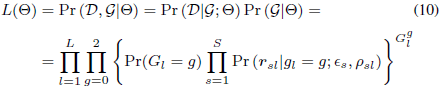

where 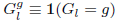 are binary indicator variables and Θ represents the set of all parameters of the model. In log-likelihood form we have:

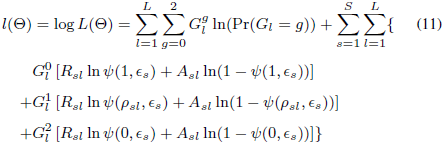

During the genotyping step, in order to maintain identifiability of the model, we fix *ρ_sl_* = 0.5 for all loci. Although *M_s_* could also be estimated within the EM procedure, we only consider overdispersion on the ASE step. These two choices lead to a much simpler EM procedure and a slightly conservative estimate of *∊_s_*.

**E-Step**: From the complete likelihood function (11) we derive the expected values for the unknown genotype indicator variables 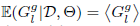 given the observed data and the current estimates for the model parameters. These quantities are also of interest for genotyping because they represent the posterior probabilities of each genotype given the data, 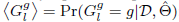:

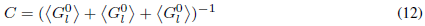

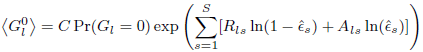

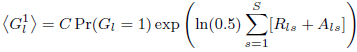

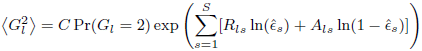

The prior genotype probabilities *Pr*(*Gl* = *g*) are obtained from the 1KG allele frequencies assuming Hardy-Weinberg equilibrium (HWE), but the user can change this.

**M-Step**: Using the expected values from the E-step, the complete likelihood is now a function of *∊s* that is easily maximized

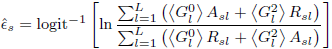

After we run QuASAR to infer genotypes across samples from the same individual, for each site we have a posterior probability of each genotype 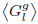, and a base-calling error 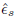 estimated for each sample. From these posteriors, discrete genotypes are called by using the genotype with the highest posterior probability; the maximum a posteriori (MAP) estimate.

**ASE-Inference:** To detect ASE we only consider sites with an heterozygous MAP higher than a given threshold (e.g., 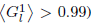). We then test the possibility that *ρ_sl_* deviates from 0.5 while also taking into account overdispersion using a beta-binomial model (combining (7) and (8)):

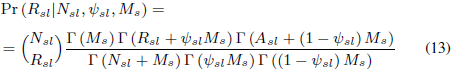

where *M_s_* controls the effective number of samples supporting the prior belief that *ρ* = .5 and is estimated using grid search:

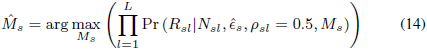

We estimate 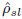 using (13) with 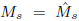 from (14) and a standard gradient method (L-BFGS-B) to maximize the log-likelihood function

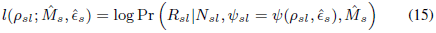

Finally, all parameters are used to calculate the LRT statistic in (9) and its *p*-value.

Additionally, we can provide an estimate of the standard error associated with the parameter *ρ_sl_* using the second derivative of the log-likelihood function (15):

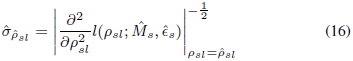

alternatively we can also recover a standard error from (9) (as is asymptotically distributed as 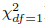), by using the *p*-value to back solve for the standard error:

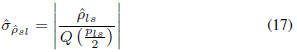

where *Q()* is the quantile function for a standard normal distribution and *p_ls_* is the *p*-value from (9). We use the first form (16) when 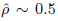 and (17) otherwise, as each provides a better approximation at those ranges. Alternatively, if we do not need 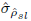 we can use (15) to obtain a profile likelihood confidence interval for *ρ_sl_*

### 2.3 Experimental Data

Lymphoblastoid cell lines (LCLs: GM18507 and GM18508) were purchased from Coriell Cell Repository and human umbilical vein endothelial cells (HUVECs) from Lonza. LCLs were cultured and starved according to (Maranville *et al.,* 2011). Cryopreserved HUVECs were thawed and cultured according to the manufacturer protocol (Lonza), with the exception that 48 hour prior to collection the medium was changed to a starvation medium, composed of phenol-red free EGM-2, without Hydrocortisone and supplemented with 2% charcoal stripped-FBS. Cells were washed 2X using ice cold PBS, lysed on the plate, using Lysis/Binding Buffer (Ambion), and frozen at -80C. mRNA was isolated using the Ambion Dynabeads mRNA Direct kit (Life Technologies). We then prepared libraries for Illumina sequencing using a modified version of the NEBNext Ultra Directional RNA-seq Library preparation kit (New England Biolabs). Briefly, each mRNA sample was fragmented (300 nt) and converted to double-stranded cDNA, onto which we ligated barcoded DNA adapters (NEXTflex-96 RNA-seq Barcodes, BIOO Scientific). Double-sided SPRI size selection (SPRISelect Beads, Beckman Coulter) was performed to select 350-500 bp fragments. The libraries were then amplified by PCR and pooled for sequencing on the Illumina HiSeq 2500 at the University of Chicago Genomics Core. For each LCL sample, libraries from 9 replicates were pooled for a total of 42.3M and 34.9M 50bp PE reads, respectively. We subsequently resequenced this libraries for a total of 81.0M and 78.6M 150bp PE reads. For the HUVEC samples (267M reads total), we collected data for 18 replicates across 6 time points to capture a wide range of basal physiologic conditions.

### 2.4 Pre-Processing

To create a core set of SNPs for ASE analysis, we first removed rare (MAF < 5%) variants from all 1KG SNPs (see Supplement for an analysis without this step). We also removed SNPs within 25 bases upstream or downstream of another SNP or short InDels as well as SNPs in regions of annotated copy numbers or other blacklisted regions (Degner *et al.,* 2012). Reads were aligned to the reference human genome hg19 using bwa mem (Li and Durbin, 2009 http://bio-bwa.sourceforge.net). Reads with quality <10 and duplicate reads were removed using samtools rmdup (http://github.com/samtools). Using a mappability filter (Degner *et al.,* 2012), we removed reads that do not map uniquely to the reference genome and to alternate genome sequences built considering all 1KG variants. Aligned reads were then piled up on the core set of SNPs using samtools mpileup command. Reads with a SNP at the beginning or at the end of the read, or with indels were also removed to avoid any potential experimental bias. Finally, the pileups were re-formatted so that each SNP has a count for reads containing the reference allele, and a count for those containing the alternate allele.

### 2.5 Comparison with Other Methods

To assess QuASAR genotyping quality, we focused on the LCL data and compared genotype calls to those of 1KG project. We also compared QuASAR genotyping accuracy to samtools + bcftools (Li and Durbin, 2009) and GATK (DePristo *et al.,* 2011). To assess QuASAR ASE inference, we used QQ-plots and eQTL derived from the GEUVADIS dataset (Lappalainen *et al.,* 2013; Wen *et al.,* 2014). When we compared QuASAR ASE inference to other methods, we used our implementation of the Binomial and Beta-binomial test by forcing QuASAR to ignore the genotyping uncertainty in the ASE inference step; i.e., in (9) numerator we only consider *ρ_sl_* =; 0.5. For the Binomial test we further force *M_s_* = ∞>. In the supplement, we also compared QuASAR performance to the method used in (Kukurba *et al.,* 2014). To make the results comparable, we used QuASAR genotype information and the same pre-processing steps described aboveupto `samtools mpileup`.

## 3 Results

We implemented the QuASAR approach as detailed in the Methods section in an R package available at http://github.com/piquelab/QuASAR. First, to evaluate QuASAR genotyping accuracy, we collected RNA-seq data for two lymphoblastoid celllines (LCLs) that already have high quality genotype calls from the 1KG project (GM18507 and GM18508). As illustrated in Table 1, we are able to accurately genotype thousands of loci with lower error rates than other methods commonly used for genotyping DNA-seq data. By design, QuASAR is more conservative in making heterozygous calls than homozygous calls, yet still retains a large number of heterozygous loci compared to GATK (DePristo *et al*., 2011) and samtools (Li and Durbin, 2009). In QuASAR, if there is contention between: i) an heterozygous genotype call with extreme allelic imbalance, or ii) an homozygous genotype call with base call errors; our model will favor the latter ii). This is a crucial feature for accurate inference of ASE, which will be discussed later in more detail. As we increase the threshold on the QuASAR posterior probability of heterozygosity we consistently reduce the genotype error rate Table S1. The error rates increase slightly if we include rare variants with MAF < 0.05 (Table S2), and are minimally affected when 1KG allele frequencies are not used as priors (Table S3). An important feature of QuASAR, is that the genotying information is also used for the next inference step.

**Table 1.** **QuASAR accuracy in genotyping heterozygous loci compared to GATK and samtools**. For four samples with different sequencing depth and read length are compared. Each row reports the number of heterozygous SNPs identified by each method and the percentage of false discoveries when compared to 1KG genotypes. For GATK and samtools we used the default parameter settings, and for QuASAR we considered SNPs with a posterior probability of heterozygosity > 0.99

| Sample<br>(# reads, read length) |  | Genotyping method |  |  |
| --- | --- | --- | --- | --- |
|  |  | Samtools | GATK | QuASAR |
| NA18507<br>($4.23 \times 10^7$, 50bp) | # hets | 3158 | 4568 | 3310 |
|  | FDR | 1.20% | 1.31% | 0.91% |
| NA18508<br>($3.49 \times 10^7$, 50bp) | # hets | 2718 | 3933 | 2844 |
|  | FDR | 0.63% | 0.97% | 0.56% |
| NA18507<br>($8.10 \times 10^7$, 150bp) | # hets | 19515 | 24522 | 17434 |
|  | FDR | 1.29% | 2.09% | 1.14% |
| NA18508<br>($7.86 \times 10^7$, 150bp) | # hets | 16998 | 21587 | 15526 |
|  | FDR | 1.08% | 1.75% | 1.08% |

We next sought to characterize QuASAR performance in genotyping and ASE-inference from RNA-seq experiments sequenced at different depths. In total we analyzed 18 samples (3 replicates across 6 different time-points) for an individual for which genotypes were not previously available. We combined the 18 fastq files in different ways as input for QuASAR, to obtain an empirical power curve (see Figure 2). As expected, we observed that our ability to detect heterozygotes (MAP> 0.99) increases with the sequencing depth. At a more modest sequencing depth of 16 million, we can still detect more than 1,000 heterozygous sites.

After obtaining the genotypes, we assessed whether there is evidence of ASE at any of the SNPs determined to be heterozygous. To conduct ASE inference we used the LRT (9) statistic in QuASAR and obtained p-values. We controlled the FDR using the q-value procedure (Storey, 2002). As shown in Figure 3, our ability to detect ASE greatly increased with the number of SNPs we were able to genotype, which in turn is a function of coverage (Figure 2). Using 100 million reads we detected roughly 9,000 heterozygous SNPs of which 50 have ASE at 10%FDR.

**Fig. 2.**
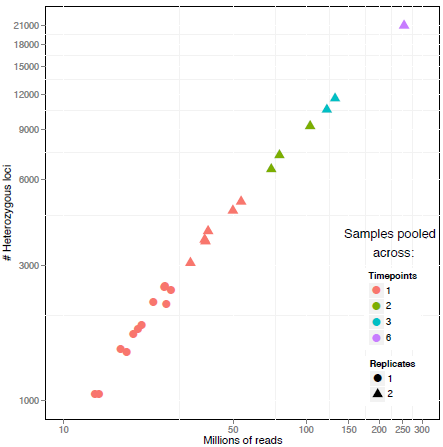
Empirical power in detecting heterozygous SNPs as a function of sequencing depth. Each point represents a single input dataset to QuASAR: either as a single experiment replicate and time point (red dot), combining multiple time points (2 = green, 3 = blue, 6 = purple), or combining replicates (1 = dot, 2 = triangle). The *x*-axis represents the total number of RNA-seq reads in the fastq input files. The *y*-axis represents the log_10_ of the total number of SNPs that are determined to be heterozygous.

To assess the calibration of the ASE test in QuASAR and to compare it to other ASE inference approaches, we used QQ-plots, Figure 4. A QQ-plot compares the quantiles observed from a test statistic to those that are expected under a null distribution (e.g., *p*-values are uniformly distributed between 0 and 1). The shape of the QQ-plot curve is useful to judge how well the *p*-values are calibrated when we expect that a large number of the tests conducted are sampled from the null distribution. In this latter scenario, we expect that the QQ-plot curve would follow the 1:1 line for the range of *p*-values with higher value. For small *p*-values, we expect that the curve starts to depart from the 1:1 line representing the small proportion of tests that are not sampled from the null distribution. Many existing approaches for ASE use either a Binomial test (McDaniell *et al.,* 2010; Reddy *et al.,* 2012; Degner *et al.,* 2012; Kukurba *et al.,* 2014) or a beta-Binomialtest (Sun, 2012; Pickrell *et al.,* 2010) that does not account for genotyping or base-calling error. To compare QuASAR ASE inference to these alternative approaches, we used the RNA-seq data collected on the LCL samples and the genotype calls from QuASAR. Figure 4 clearly shows that the Binomial test is too optimistic, and will likely lead to many false discoveries. Another Binomial test (Figure S1) independently implemented (Kukurba *et al.,* 2014) show a similar behaviour to our own implementation. In contrast to the Binomial test, the Beta-binomial model seems better calibrated, but uncertainty on the genotype being a true heterozygote can lead to very small *p*-values and false positives. QuASAR combines the Beta-binomial model with uncertainty on the genotype, resulting in the most conservative approach, likely avoiding a common cause of false positives in ASE analysis.

**Fig. 3.**
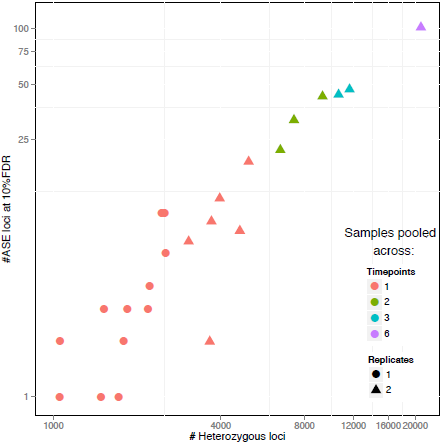
Empirical power in detecting ASE as a function of the number of heterozygous SNPs detected. Each point represents a single input dataset to QuASAR as in Figure 2. The *x*-axis represents the total number of SNPs that are determined to be heterozygous. The *y*-axis represents the log_10_ of the number of SNPs that have a significant *p*-value for ASE at 10% FDR.

To further evaluate the differences between QuASAR and other methods, we focused on the high coverage LCL samples. This cell-type has been part of large sample eQTL studies such as GEUVADIS (Lappalainen *et al.,* 2013) including RNA-seq data for more than 600 individuals. From a recent reanalysis of the GEUVADIS dataset (Wen *et al.,* 2014) we selected genes with eQTLs (5% FDR) and a leading SNP with a high posterior inclusion probability (PIP> 0.9), as those are the nucleotides more likely to be causal (i.e., the eQTN). We hypothesized that in our data, genes for which the eQTN was heterozygote would be more likely to show evidence for ASE than those for which the eQTN was homozygote. However, transcripts with homozygous eQTNs could still show ASE, if they have additional personal/rare associations or due to an imprinting mechanism. In any of the methods we tested (Figure S2), there was a significant difference between ASE *p*-values in the expected direction (Mann-Whitney U-test *p* < 0.02). The distribution of p-values (Figure S2 D) for ASE signals in transcripts with homozygous eQTNs are much closer to the uniform distribution in QuASAR when compared to other methods, especially those that do not account for overdispersion.

In general, the *p*-values obtained across all methods tested are very much correlated to each other, as shown in Figure S3 but there are key differences. The methods that use a binomial test tend to have lower *p*-values than QuASAR which uses a beta binomial model. Figure S4 A and B show that we may observe more shrinkage for SNPs with deeper coverage. Compared to a test using the beta-binomial distribution but ignoring genotype uncertainty, our QuASAR test leads to similar *p*-values. but SNPs with more uncertainty on being heterozygous are corrected in a higher degree towards a less significant *p*-value (Figure S3 C and S4 C id="fig4" position="float" fig-type="figure">

**Fig. 4.**
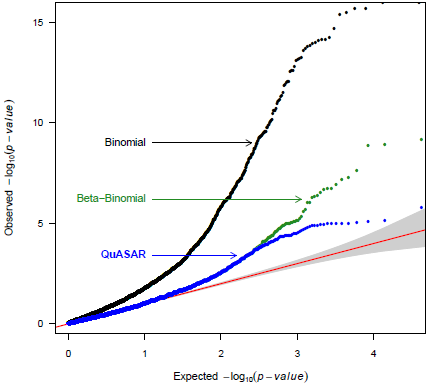
QQplot comparing the p-value distribution of 3 alternative methods for determining ASE. The *x*-axis shows the log_10_ quantiles of the p-values expected from the null distribution. The *y*-axis shows the log_10_ quantiles of the *p*-values computed from the real data using 3 different methods: i) Binomial (black) assumes *M* = ∞ no overdispersion; ii) Beta-binomial (green) considers overdispersion but does not consider uncertainty in the genotype; iii) QuASAR (blue) uses the Beta-binomial distribution and uncertainty in the genotype calls. In all three cases the same set of SNPs are considered. The shaded area in gray indicates a 95% confidence band for the null distribution.

In terms of computational complexity. QuASAR is very fast. QuASAR runtime for genotyping with the EM procedure takes less than 20 seconds on any of our data-sets (using a computer with a 6 core 2.93GHz Intel processor and 8GB RAM). Each EM iteration in the genotyping step is *O*(*LS*) linear with the number of SNPs and samples, and convergence is achieved in less than 20 iterations. GATK with the default options takes a longer amount of time (about 2 hrs.) compared to samtools and QuASAR. as the latter methods focus directly on 1KG SNPs only. Comparatively, for both our pipeline and (Kukurba *et al.,* 2014). more time is spent in data pre-processing (roughly 10 minutes per sample depending on the sequencing depth and assuming reads are already aligned) than in genotyping or ASE inference.

## 4 Discussion

QuASAR is the first approach that detects genotypes and infers ASE from the same sequencing data. In this work, we focused on RNA-seq. but QuASAR can be applied to other data types (ChIP-seq, DNase-seq, ATAC-seq. and others). Indeed, the more experimental data available from the same individual across many experimental samples (data-types, conditions, cell-types or technical replicates), the more certainty we can gather about the genotype.

A key aspect of the QuASAR ASE inference step is that it takes into account over-dispersion and genotype uncertainty resulting in a test that we have shown to be well-calibrated. In many cases, the p-values obtained from biased statistics can be recalibrated to the true null distribution using a permutation procedure. Unfortunately, this is not possible for ASE inference, as randomly permuting the reads assigned to each allele would inadvertently assume that the reads are distributed according to a Binomial distribution. More complicated and computationally costly resampling procedures can be proposed, but it is not clear what additional assumptions may be introduced and if such methods can correctly take into account genotyping uncertainty.

If prior genotype information is available, it can also be provided as input to the algorithm. The prior uncertainty of the genotypes should be reflected in the form of prior probabilities for each genotype. In this paper, we have shown that we can obtain reliable genotype information from RNA-seq reads, thus making additional genotyping unnecessary if the endpoint is to detect ASE. Instead, sequencing the RNA-seq libraries at a higher depth is probably a better strategy as it greatly improves the power to detect ASE.

Furthermore, as sequencing costs decrease rapidly. ASE methods are becoming very attractive in applications where eQTL studies have been previously used. This is of increased importance in scenarios where collecting a large number of samples is expensive or infeasible. Large scale eQTL studies are still very much necessary for fine-mapping, but analysis of ASE can provide unique insights into mechanisms that are uncovered only under specific experimental conditions, for example as a result of gene x environment interactions.

## Acknowledgement

We would like to thank Wayne State University HPC Grid for computational resources, the University of Chicago Genomics Core for sequencing services, and Jacob Degner and members of the Luca/Pique group for helpful comments and discussions.

Funding: NIH 1R01GM109215-01 (RPR, FL)AHA 14SDG20450118 (FL)

## Supplementary Information

### Supplementary Figures

**Table S1.** **QuASAR accuracy in genotyping**. Each row reports the number of heterozygous and homozygous SNPs identified by QuASAR and the percentage of false discoveries when compared to 1KG genotypes. Each column uses a different MAP threshold to define high confidence genotype calls.

| Sample | QuASAR Performance | MAP $\langle G_l^g \rangle$ | | |
| --- | --- | --- | --- | --- |
|  |  | >0.5 | >0.9 | >0.99 |
| NA18507 | Heterozygotes | 3320 | 3316 | **3310** |
|  | False Discoveries | 0.93% | 0.93% | 0.91% |
|  | Homozygotes | 9989 | 9984 | 9966 |
|  | False Discoveries | 3.94% | 3.92% | 3.90% |
| NA18508 | Heterozygotes | 2858 | 2851 | **2844** |
|  | False Discoveries | 0.59% | 0.60% | 0.56% |
|  | Homozygotes | 8598 | 8594 | 8576 |
|  | False Discoveries | 4.06% | 4.01% | 3.99% |

**Table S2.** **QuASAR accuracy in genotyping when rare alleles are included**. Rows and columns have the same meaning as in Table S1 but we have not excluded genetic variants with MAF< 0.02. These results use the 1KG allele frequencies and HWE to calculate the genotype prior probability.

| Sample | QuASAR Performance | MAP |  |  |
| --- | --- | --- | --- | --- |
|  |  | 0.5 | 0.9 | 0.99 |
| NA18507 | # heterozygotes | 3289 | 3282 | 3276 |
|  | FDR | 2.77% | 2.65% | 2.59% |
|  | # homozygotes | 89209 | 89203 | 89184 |
|  | FDR | 0.84% | 0.84% | 0.84% |
| NA18508 | # heterozygotes | 2871 | 2867 | 2861 |
|  | FDR | 1.32% | 1.22% | 1.15% |
|  | # homozygotes | 76835 | 76830 | 76816 |
|  | FDR | 0.76% | 0.76% | 0.75% |

**Table S3.**
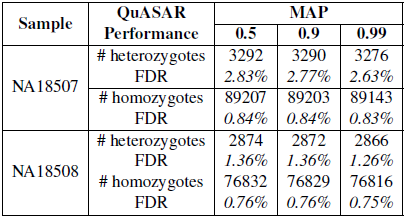
**QuASAR accuracy in genotyping when rare alleles are included and 1KG allele frequencies are not used as prior information.** Rows and columns have the same meaning as in Table S2, and we also have not excluded genetic variants with MAF< 0.02. These results do not use 1KG allele frequencies for calculating the genotype prior probability, MAF= 0.5 and HWE is used for all SNPs.

**Fig. S1.**
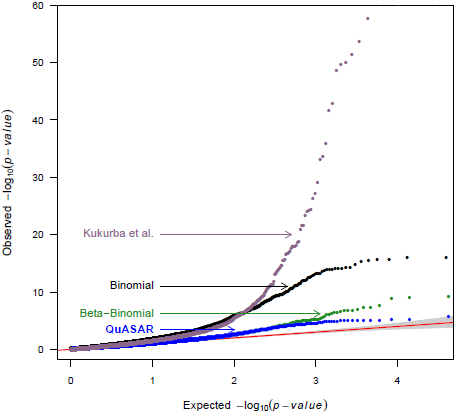
QQplot comparing the p-value distribution of QuASAR to alternative methods for determining ASE. Three of the methods are the same as in Figure 4: i) Binomial (black), ii) Beta-binomial (green), iii) QuASAR (blue), and another iv) Binomial method as implemented by (Kukurba *et al,* 2014). In all cases the same set of SNPs and genotypes are considered, but iv) uses a different pre-processing pipeline that may filter less reads. The shaded area in gray indicates a 95% confidence band for the null distribution.

**Fig. S2.**
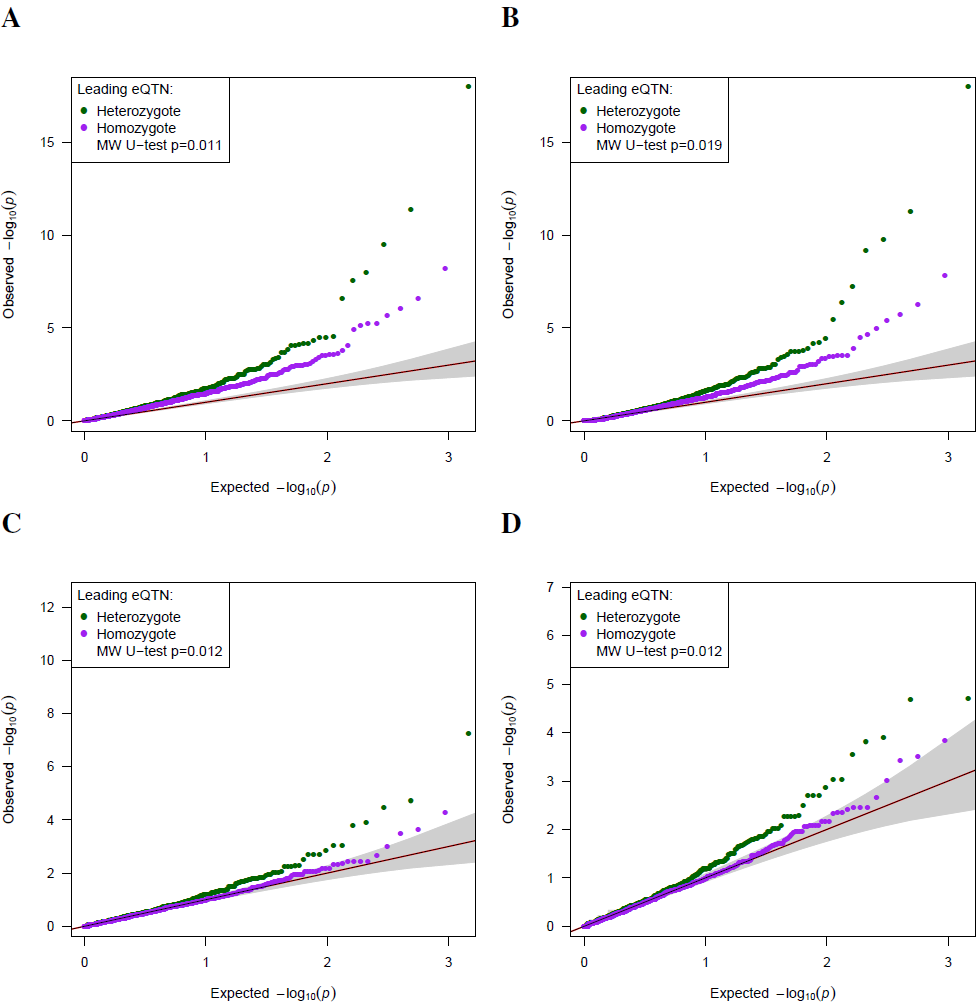
Qplot comparing the ASE *p*-value distribution for transcripts with known eQTLs in LCLs. Each panel corresponds to one method: **A)** Binomial (our implementation), B) Binomial (Kukurba *et al,* 2014), **C)** Beta-binomial (our implementation), D) QuASAR (our proposed method). Each dot corresponds to a heterozygous SNP in a gene transcript for which an eQTL is previously known in LCLs. Only eQTLs with a leading SNP having a high posterior inclusion probability (PIP> 0.9 as defined by Wen *et al,* 2014) are used for this analysis and denoted as eQTN. The ASE measurements are then divided in two groups: (green dots) those with an eQTN that has a heterozygous genotype and are expected to show some evidence of ASE, and (purple dots) those with an eQTN with a heterozygous genotype and are less likely to show evidence for ASE. The legend also reports the Mann-Whitney U-test comparing whether the green-dots tend to have a lower p-value than the purple-dots. The shaded area in gray indicates a 95% confidence band for the null distribution.

**Fig. S3.**
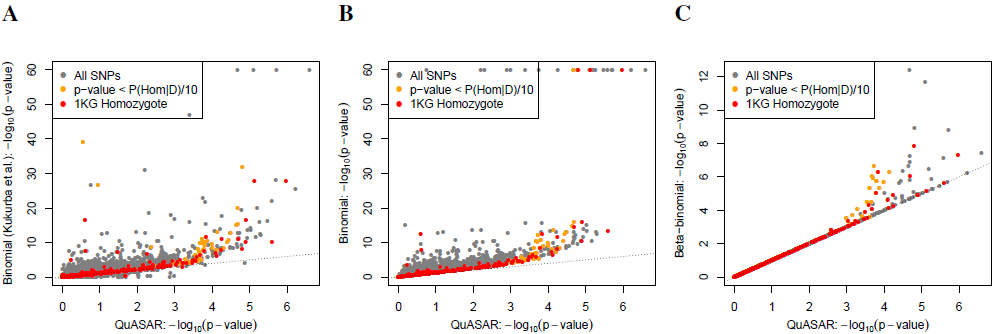
Scatter plots comparing the p-values obtained with different ASE methods. The *x*-axis represents the *—log*^10^ (*p*-values) obtained from QuASAR, and the *y*-axis the *—log*^10^ (*p*-values) for the following methods: A) Binomial (Kukurba *et al.*, 2014), B) Binomial (our implementation), and C) Beta-binomial (our implementation). Highlighted in red are those SNPs in which the genotypes from 1KG are not heterozygous. Highlighted in orange are SNPs for which the posterior probability of not being heterozygous is 10 times higher than the ASE p-value.

**Fig. S4.**
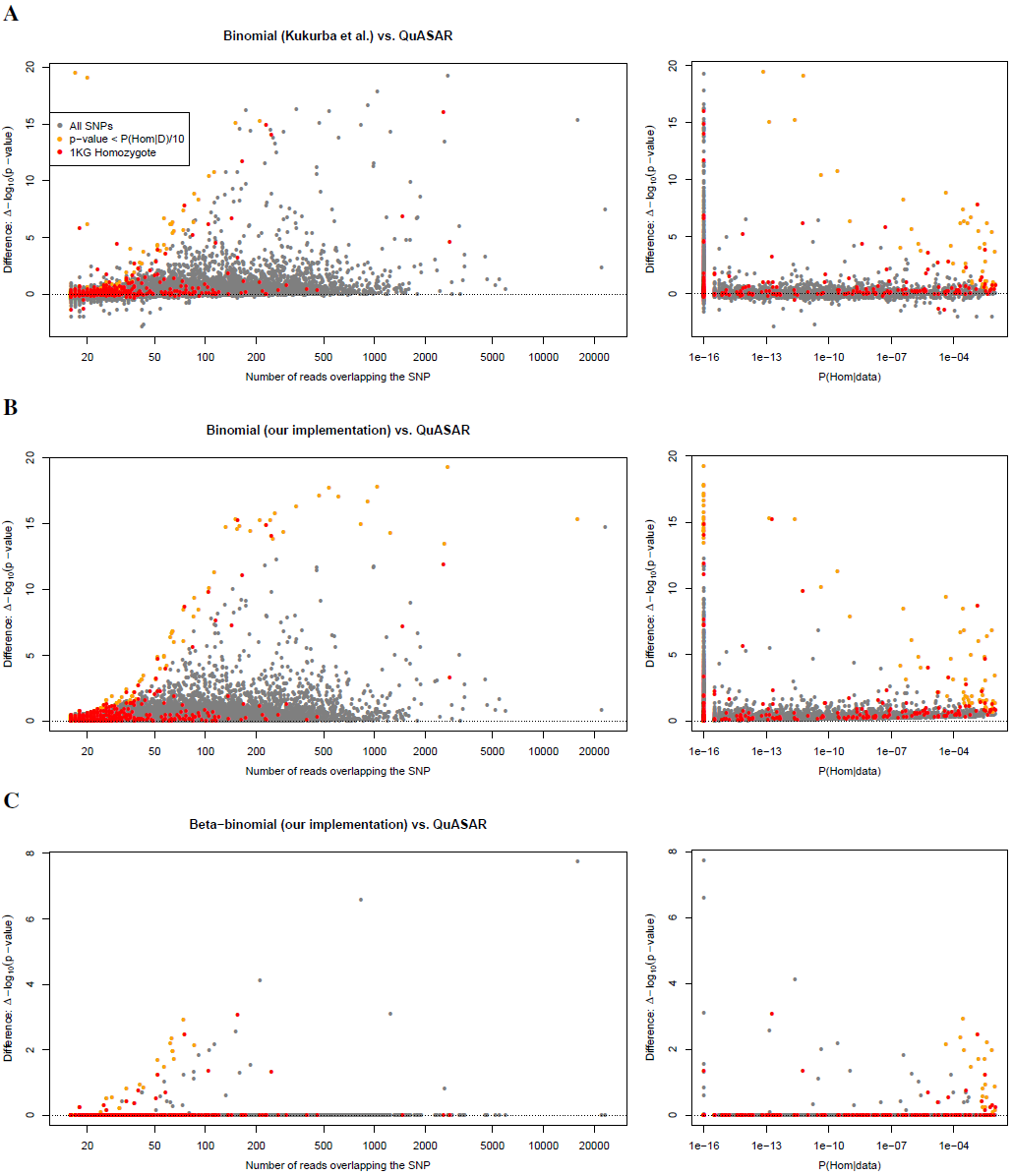
Scatter plots comparing the ASE p-values obtained with different methods as a function of read depth and probability of miscalling heterozygosity. The *y*-axis represents the difference between *—log*^10^ (*p*-values) obtained from QuASAR and those from the following methods: A) Binomial (Kukurba *et al,* 2014), B) Binomial (our implementation), and C) Beta-binomial (our implementation). The *x*-axis represents: (left) the number of reads covering that SNP in our pipeline, and (right) the QuASAR posterior probability of making a homozygous call given the data. Highlighted in red are those SNPs in which the genotypes from 1KG are not heterozygous. Highlighted in orange are SNPs for which the posterior probability of not being heterozygous is higher than 10 times the ASE p-value.

